# Environmental Impacts of Producing Culture Medium Consisting of Serum-free, Food and Complex Ingredients for Cultivated Meat

**DOI:** 10.1101/2024.09.05.611339

**Authors:** Natsufumi Takenaka, Kimiko Hong-Mitsui, Kazuhiro Kunimasa, Kotaro Kawajiri, Chihiro Kayo, Naoki Yoshikawa

**Author notes:** Corresponding Author: **Natsufumi Takenaka_****, Naoki Yoshikawa_**.

## Abstract

Culture medium accounts for most of the environmental impact of cultivated meat production. This study quantitatively evaluated and analyzed the environmental impact of producing a culture medium consisting of serum-free, food and complex ingredients for cultivated meat by performing a life cycle assessment (LCA) based on activity data at the laboratory scale. In addition, specific measures were proposed to reduce the environmental impact further. LCAs were performed at current and future production scales. This study also evaluated the impact of multiple electricity sources on the environmental impact of culture medium production. Energy, animal-derived materials, and expendables at the current scale, and energy, animal-derived materials, amino acids, and glucose at future scale are hotspots in the environmental impact of this culture medium production. The production of serum substitutes accounts for most of the environmental impact. As the scale shifts, the environmental impacts are expected to decrease by more than 70% in all impact categories. As the composition of electricity sources changed, the impact on certain categories decreased. However, as the share of renewable energy increased, the impact on land use also increased significantly. This study promotes the practical application of new culture-media for low-cost and low-environment-impact cultivated meat.

Graphical abstract.
SYNOPSIS. This study evaluated the environmental impact of culture-medium production for cultivated meat and reports that energy, animal-derived materials, expendables, amino acids and glucose are hotspots of this impact.

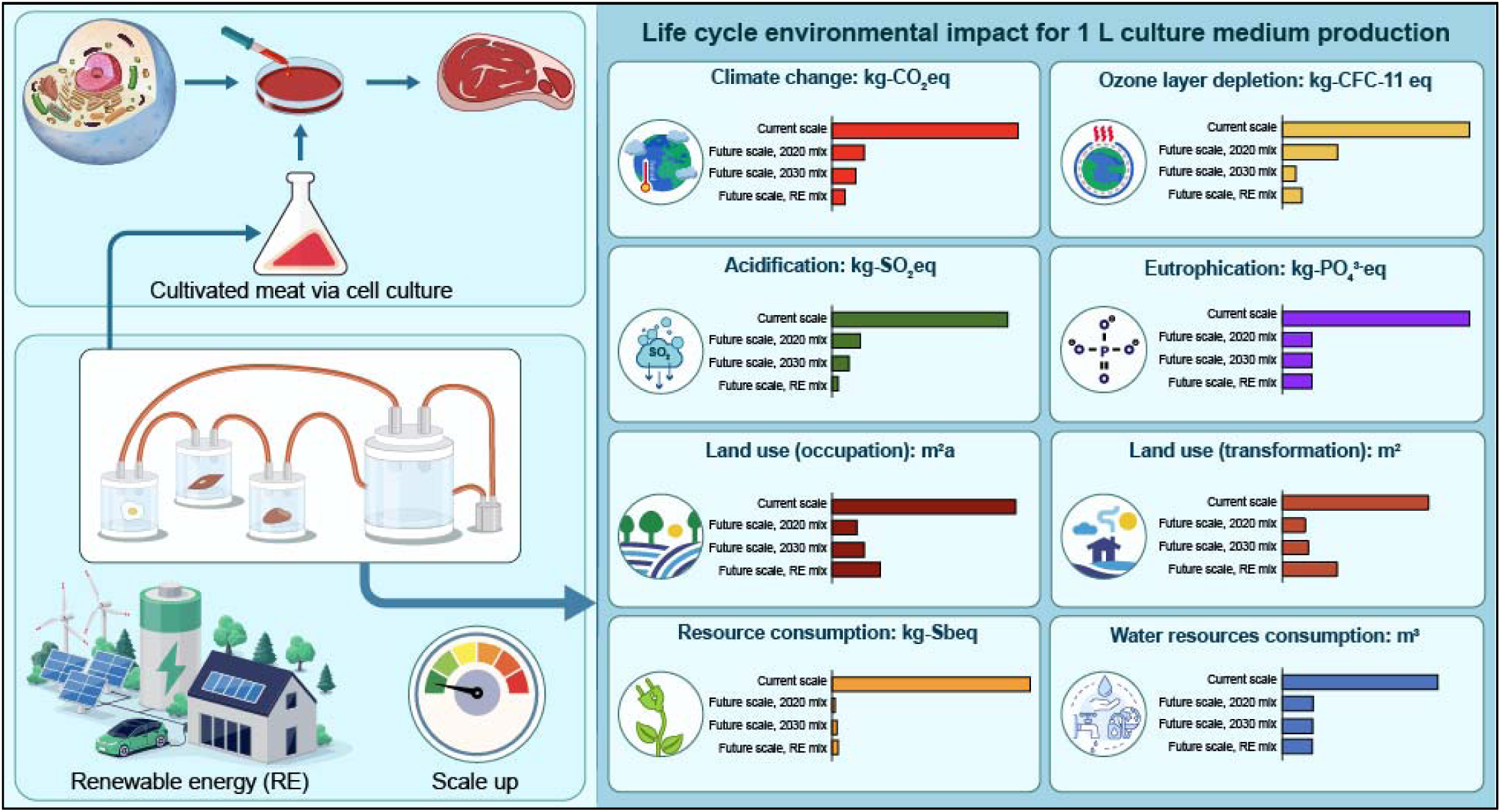

## 1. INTRODUCTION

Current meat production from livestock farming is one of the major sources of anthropogenic environmental impacts.^1^ Consuming animal-based foods has a larger environmental impact than other foods, with respect to greenhouse gas emissions, land use, water use, eutrophication, and biodiversity.^2^ Additionally, meat consumption will increase with human population growth and affluence increases, accordingly global meat production is expected to increase by 76% between the beginning of this century and 2050.^1,3^ Therefore, the environmental impact of livestock production will further increase.

Cultivated meat has attracted considerable attention as a new method of meat production. Cultivated meat aims to replicate conventionally produced meat via (stem) cell and tissue culture.^4^ It is considered to have a low environmental impact and is expected to be an alternative meat that contributes to food security in the future.^5,6^ Previous studies on the life cycle assessment (LCA) of cultivated meat have shown that environmental impacts of culture medium account for 48-89% of the total environmental impacts of cultivated meat production depending on the impact category,^7^ thus, the culture medium is a hotspot for the environmental impacts of cultivated meat production.^7,8^ Therefore, research focusing on the environmental impact of culture medium is essential for reducing the environmental impact of cultivated meat production. Previous studies on the environmental impact assessment of cultivated meat production have focused on prospective future production systems where the production technology is developed on an industrial scale, and no studies have been conducted on current production systems.^9,10^ Consequently, there is a problem that only optimistic and prospective evaluation results get ahead of themselves, and the actual environmental impact is not understood. Therefore, it is important to recognize the current situation and bridge the gap between it and the future scale.

A culture medium is a liquid that contains nutrients required for cell culture, such as carbohydrates, proteins, growth factors, minerals, and vitamins.^5^ Currently, culture medium containing fetal bovine serum (FBS) is widely used as it contains natural embryonic growth-promoting factors for enhancing cell growth and proliferation. ^11–13^ However, FBS has challenges such as price instability and the possibility of containing toxic substances such as endotoxins.^13–15^ It has also been shown that FBS contributes the most to the environmental impact of culture medium production.^11^ Therefore, a shift to a serum-free culture medium is required for cultivated meat production.^16^ Additionally, many culture media use chemical reagents. However, using ingredients not approved as food in edible cultivated meat production presents complex regulatory challenges, which are both time-intensive and expensive.^17^ Therefore, culture medium needs to be consisted of food ingredients to develop and sell edible cultivated meat products.^18^ Culture media are classified into chemically defined media, in which the components and concentrations are defined, and complex media, in which the components and concentrations are not defined as they include undefined components of natural origin such as yeast extract, meat extract, peptone, casein hydrolysate, carcass meal, or plant seed flour.^18–20^ In general, complex medium ingredients is rich in nutrients, and their use effectively reduces the price of the culture medium.^18^ Therefore, focusing on culture media consisting of serum-free, food and complex ingredients is essential for the future development of cultivated meat.

Some studies have been published about the environmental impact assessment of the culture medium production for cultivated meat. Wali et al. 2024 ^11^ performed LCA of culture medium containing FBS and alternative FBS-free media containing serum substitutes, protein hydrolysates, and recombinant growth factors based on the data from literature and published protocols. Nikkhah et al. 2024 ^21^ performed LCA on animal component-free culture media production using recombinant growth factors and proteins as serum substitute components by modeling systems on a theoretical, commercial scale. However, no previous studies have assessed the environmental impact of producing culture medium that consists of serum-free, food and complex ingredients, nor have they conducted LCA based on the company’s actual culture medium production data.

Based on the above background, this study conducted an LCA based on a company’s actual culture medium production data to evaluate the current environmental impact of culture medium consisting of serum-free, food and complex ingredients for cultivated meat production. This study also conducted the LCA contingent on this culture medium production being more established and on a larger scale in the future and using different electric power supply configurations. Furthermore, effective strategies for reducing the environmental impact of culture medium production were examined based on the LCA results. The significance of this study is threefold. First, this study quantitatively evaluated the culture medium’s current and future environmental impacts, a key technology for low-cost and low-environmental impact on cultivated meat production. Second, this study identified hotspots in the environmental impact of culture medium production. Third, this study proposes specific measures to further reduce the environmental impact. These results support the practical application of new culture media for low-cost and low-environment-impact cultivated meat.

## 2. MATERIALS AND METHODS

This study used LCA for environmental impact assessment. LCA is a method for quantitatively assessing the environmental impact over the life cycle of a product or service and is standardized by the International Organization for Standardization (ISO).^22,23^

### 2.1 Functional Unit and System Boundary

The functional unit of this study was 1 L of culture medium. The system boundary of this study was cradle-to-gate, from raw material production to product manufacturing. Figure 1 shows a schematic of the system boundary used in this study. Sterilization of culture medium, device assembly, autoclaving, and gas sterilization of bioreactors, devices, and equipment are included in the evaluation scope of this study. The disposal of waste generated in each production process and each device’s cleaning process after use were also included. As the environmental impact of equipment manufacturing is small compared with that of equipment operation when the lifetime is considered, manufacturing the main body of the device and the plant was not included in this study’s evaluation scope.^8^

**Figure 1.**
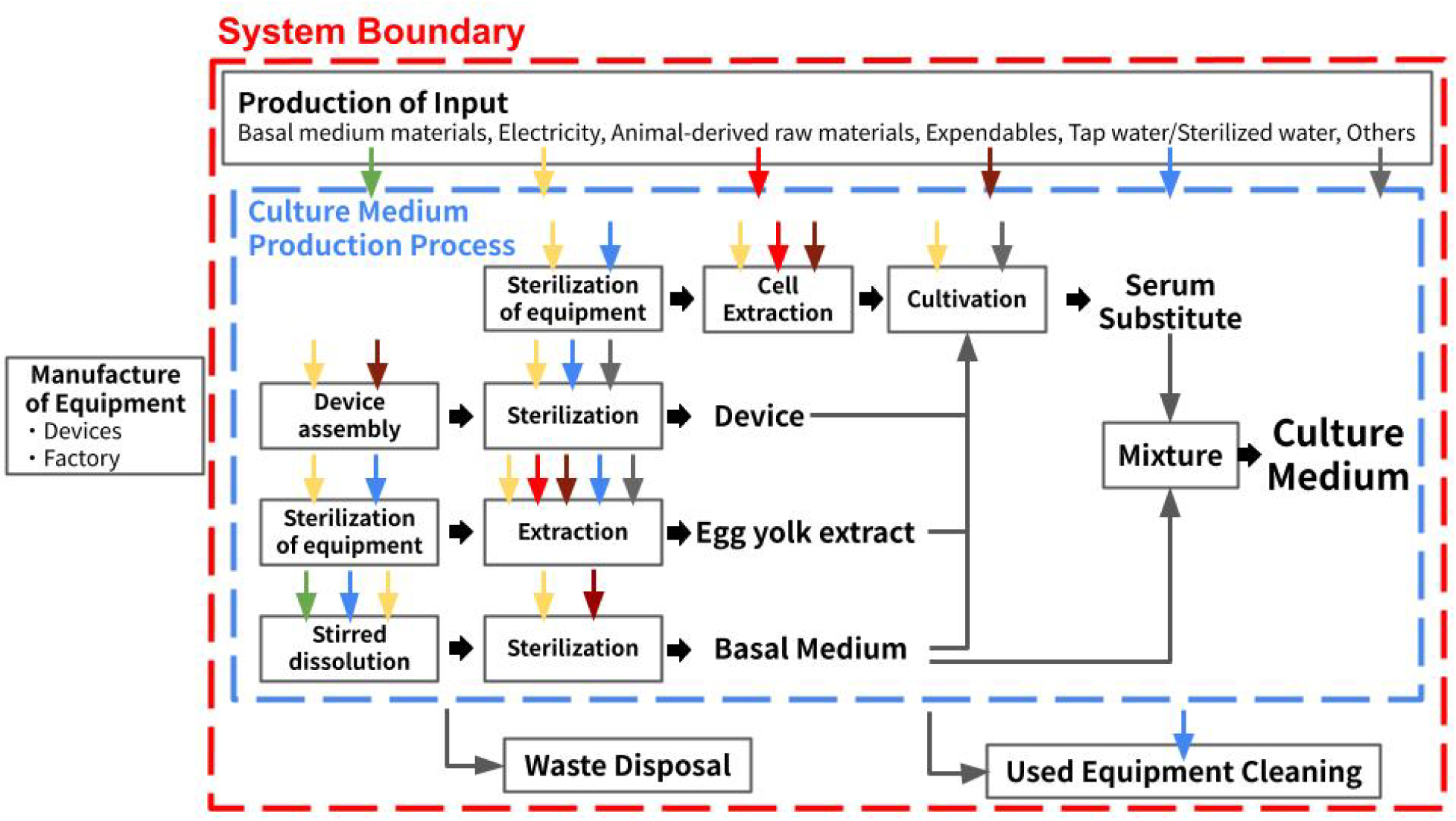
System boundary schematic. The red frame indicates the system boundary, the blue frame indicates the production process of this study’s culture medium, the square box indicates each process, and the flow color corresponds to the input production. The flow for each process indicates the input used in that process.

### 2.2 Target Culture Medium

The culture medium developed by IntegriCulture Inc. (Japan) was the subject of this study. The basal medium of this study’s target culture medium was designed based on Dulbecco’s Modified Eagle’s Medium (DMEM), with all components of DMEM replaced with food/food additives.^24^ The serum substitute components were produced using a method in which multiple organ cells are co-cultured using the above basal medium, and the interaction of each organ cell secretes components that serve as a serum substitute to achieve a serum-free culture medium. The target culture medium was prepared by mixing equal amounts of the food ingredient basal medium and complex serum substitutes.

The basal medium was prepared by dissolving the raw materials in sterile water. Then, the basal medium was filter sterilized. Serum substitute components were produced by co-culturing multiple organ cells in the above basal medium. Specifically, as shown in Figure 2, the serum substitute components production system consists of one adjustment reactor without cell culture and three feeder reactors with cell culture, all of which are connected. All reactors were 0.6 L in size and filled with a total of 1.2 L of basal medium. Once a week after the start of the culture, half of the culture supernatant (0.6 L) was collected, and 0.6 L of the new basal medium was added. The period of one batch was 4 weeks, and all culture supernatants were collected in the final week.

**Figure 2.**
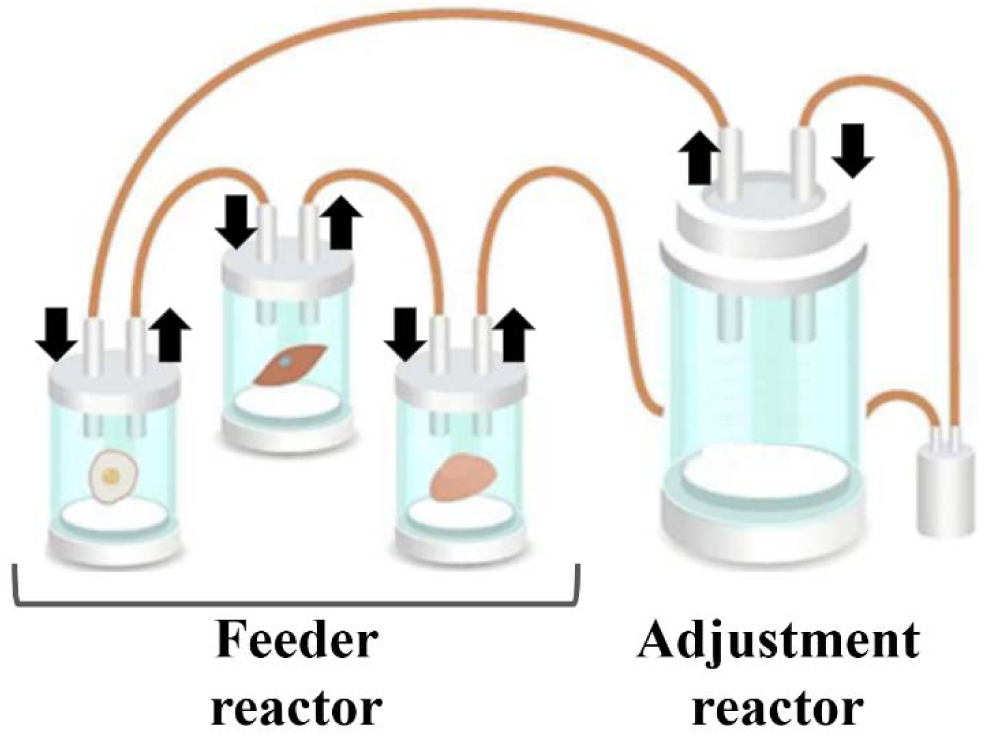
Cell culture for the preparation of serum substitutes. Figure 2 shows the cell culture methods for preparing serum substitutes. Adjustment reactor contains only a culture medium without cell culture. Three feeder reactors were used for cell culture. All reactors were connected, and the medium was circulating. The black arrows indicate the direction of circulation.

All culture supernatants collected weekly were serum substitute components for the target culture medium, produced 3 L per batch. Mixing equal amounts of the serum substitute components with the basal medium produced 6 L of culture medium per batch.

Cells were extracted from eggs. Bioreactors and devices for exchanging the culture medium in the cell culture process used silicone tubes and other experimental expenses for conducting lines. At the start-up of each batch, the bioreactor, equipment and instruments were assembled and sterilized. They were cleaned by hand at the end of the batch. Autoclaving and gas sterilization were used as sterilization methods. Egg yolk extract and yeast extract were used as culture aids in the cell culture process to prepare serum substitutes. Egg yolk extract was only used at the start of the culture, and the final proportion of the culture medium was 2%.

The cell culture ability of the basal medium used in this study is summarized in Kanayama et al. 2022 ^24^, where its alternative to DMEM is recognized. Duck liver cells were cultured for 2 days from the same initial cell number in this study’s basal medium/FBS 10% and this study’s culture medium (n=3). This study’s culture medium showed a 107% viability rate, compared with this study’s basal medium/FBS 10%. The details of this cell culture are in S.1 of the Supporting Information (SI).

### 2.3 Scenario Analysis

This study assessed the environmental impact of this culture medium production at present and future scales. The impact of multiple electricity sources on the environmental impact of culture medium production was also assessed at future scale.

#### 2.3.1 Scale

Assessing the current and actual environmental impact of culture medium production is important to bridge the gap between the current situation and future scale. As this study collected data in 2023, the assumed year for the current scale is 2023. Conversely, the current scale is the lab-scale, which implies that the process efficiency of small-scale systems tends to be very low and might not provide any practical insight into the industry. Therefore, this study conducted LCA for the case where future culture medium production is established on a larger scale. Thereby this research identifies the environmental impact of this culture medium production on a commercial scale and shows its potential. This will help to explore the directions for future development of low-environment-impact culture medium production. As a future scale, this study estimated an even larger scale for the scale-up plans of the target company. For the estimation of a 10,000 L reactor (the largest reactor assumed for use at the future scale), reference was made to Sinke et al. 2023 ^8^, a previous LCA study that predicted cultivated meat production on a commercial scale in 2030. Therefore, the assumed year for the future scale in this study is 2030.

Table 1 summarizes the changes from present to future scales. The future scale will have the same reactor configuration, cultivation method, and duration as the current scale. The size of the adjustment reactor was assumed to be 10,000 L and the size of the feeder reactor 100 L. 12,500 L of serum substitute components are produced per batch, and 25,000 L of culture medium is produced per batch. The current scale used silicone tubes and other expendables conducting lines. In the future, it is assumed that all piping from the main body of the device to the conductors will be stainless-steel piping for multiple uses. The cleaning method will assumably change from hand washing to Cleaning in Place (CIP), and the sterilization method will change from autoclaved and gas sterilization to Sterilization in Place (SIP) using steam. In the cell culture process for preparing serum substitutes, the temperature and air were regulated by placing all bioreactors in a single incubator at the current scale. However, it is assumed that the temperature and air will be controlled for each bioreactor in the future. Cell line is a general term that applies to a defined population of cells that can be maintained in culture for an extended period, retaining the stability of certain phenotypes and functions.^25^ Cell lines that divide semi-permanently are generally considered a requirement for generating huge amounts of edible tissue from a stable, robust bioprocess, as they have the advantage that they do not need to rely on animals as a source of cell extraction and can prepare cells in large quantities.^26,27^ However, particularly in Japan, consumer acceptance of using cell lines in cultivated meat production is uncertain, therefore, primary cells extracted from eggs are used in the current scale.^26,27^ Conversely, it was assumed that the development of edible cell lines, proof of the safety of cell line use, and consumer understanding would lead to the introduction of cell lines and a shift from eggs to cell lines as a source of cell extraction on a future scale.

**Table 1.**
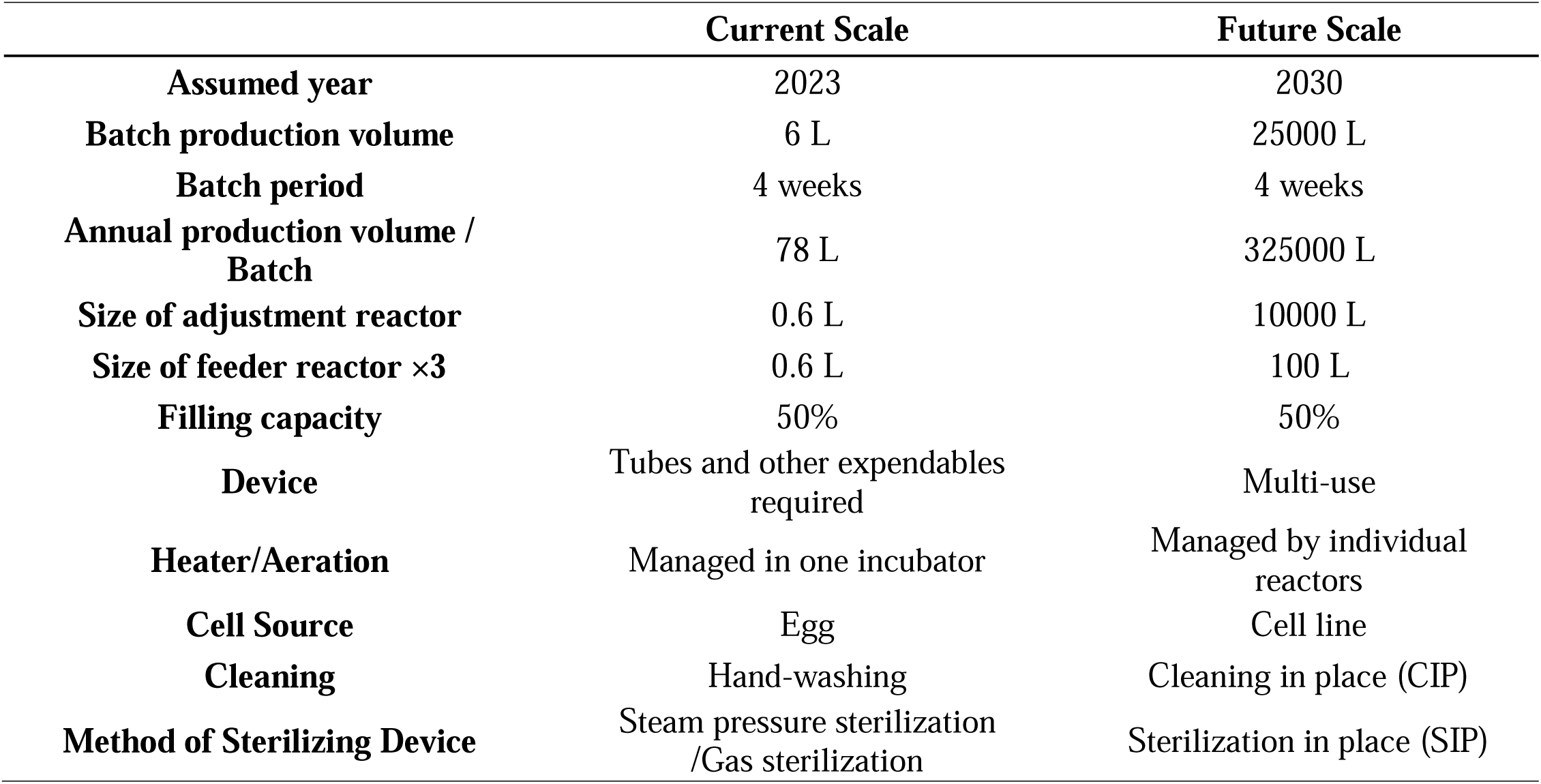
Comparison of present and future scales.

#### 2.3.2 Energy

This study used data on Japan’s average electricity source composition in 2020 (“Current Electricity Mix”) as the energy source used at the current scale because it was the latest data in the inventory database. For environmental impact assessments of the future scale, “Current Electricity Mix” was also used to clarify the differences between current and future scales excluding future changes in the electricity mix. In addition to the “Current Electricity Mix,” future-scale LCA was conducted using Japan’s expected electricity source composition in 2030 (“Electricity Mix in 2030”) announced by the Japanese Agency for Natural Resources and Energy, and electricity source composition for solar and wind power generation only (“Renewable Electricity Mix”).^28^ Table 2 shows the specific electricity source composition for “Current Electricity Mix,” “Electricity Mix in 2030,” and “Renewable Electricity Mix.”

**Table 2.** Electricity source composition of each energy mix.

|  | **Current Mix** | **Electricity Mix in 2030** | **Renewable Electricity Mix** |
| --- | --- | --- | --- |
| **Solar** | 1.9% | 16.0% | 50.0% |
| **Wind** | 0.9% | 5.0% | 50.0% |
| **Petroleum etc.** | 1.7% | 2.0% | - |
| **Coal** | 32.5% | 19.0% | - |
| **Liquefied Natural Gas (LNG)** | 41.9% | 20.0% | - |
| **Other gases** | 3.1% | - | - |
| **Nuclear** | 4.4% | 20.0% | - |
| **Geothermal** | 0.2% | 1.0% | - |
| **Hydropower** | 10.0% | 11.0% | - |
| **Biomass** | 2.3% | 5.0% | - |
| **Waste** | 0.4% | - | - |
| **Hydrogen, Ammonia** | - | 1.0% | - |
| **Others** | 0.7% | - | - |

Therefore, in this study, as shown in Table 3, LCA was performed for three scenarios in addition to the current scale, which is the actual and current culture-medium production process. All three scenarios targeted the future scale and used the following data as electricity sources: “Current Electricity Mix” for [Future Scale, 2020 mix], “Electricity Mix in 2030” for [Future Scale, 2030 mix], and “Renewable Electricity (RE) Mix” for [Future Scale, RE mix]. Regarding the use of heat, town gas-derived heat was assumed as the energy source for CIP, SIP, and initial heating of the medium at the future scale. The prediction methodology is described in S.2.3 and S.3 in SI.

**Table 3.** Scenario Description.

| Scenario | Scale | Electricity Source |
| --- | --- | --- |
| **Current Scale** | Current | Current Electricity Mix |
| **Future Scale, 2020 mix** | Future | Current Electricity Mix |
| **Future Scale, 2030 mix** | Future | Electricity Mix in 2030 |
| **Future Scale, RE mix** | Future | Renewable Electricity (RE) Mix |

### 2.4 Data Collection

IntegriCulture Inc. (Japan) provided data for the culture medium used in this study. Data were collected from the company’s standard operating procedure for culture medium production and the entire production process from March 2023 to June 2023. The data on the composition of the basal medium used in this study were obtained from the literature.^24^ The energy consumption of bioreactors and the energy and input consumption of SIP/CIP at future scales were predicted from the information in the literature. The details of the future-scale LCA methodology in this study are described in S.2 and S.3 of SI.

### 2.5 Life Cycle Inventory (LCI)

Table 4 shows inventory data for the evaluation scope. It summarizes the classifications and materials, amounts used per liter of culture medium production, data sources, and purposes for each present and future scale. Each input and output were classified by electricity, expendables, waste, amino acids, glucose, vitamins + inorganic components + trace elements, animal-derived raw materials, tap water + sterilized water, and others. The results and other information are presented based on this study’s classification. The target culture medium production used sterilization filters, pipettes, tubes, 100 mm dishes, reactor components, paper and plastic tweezers, classified as expendable. Sugar beet is the main source of amino acids, and maize is the main source of glucose, the raw material for the basal medium.

**Table 4.**
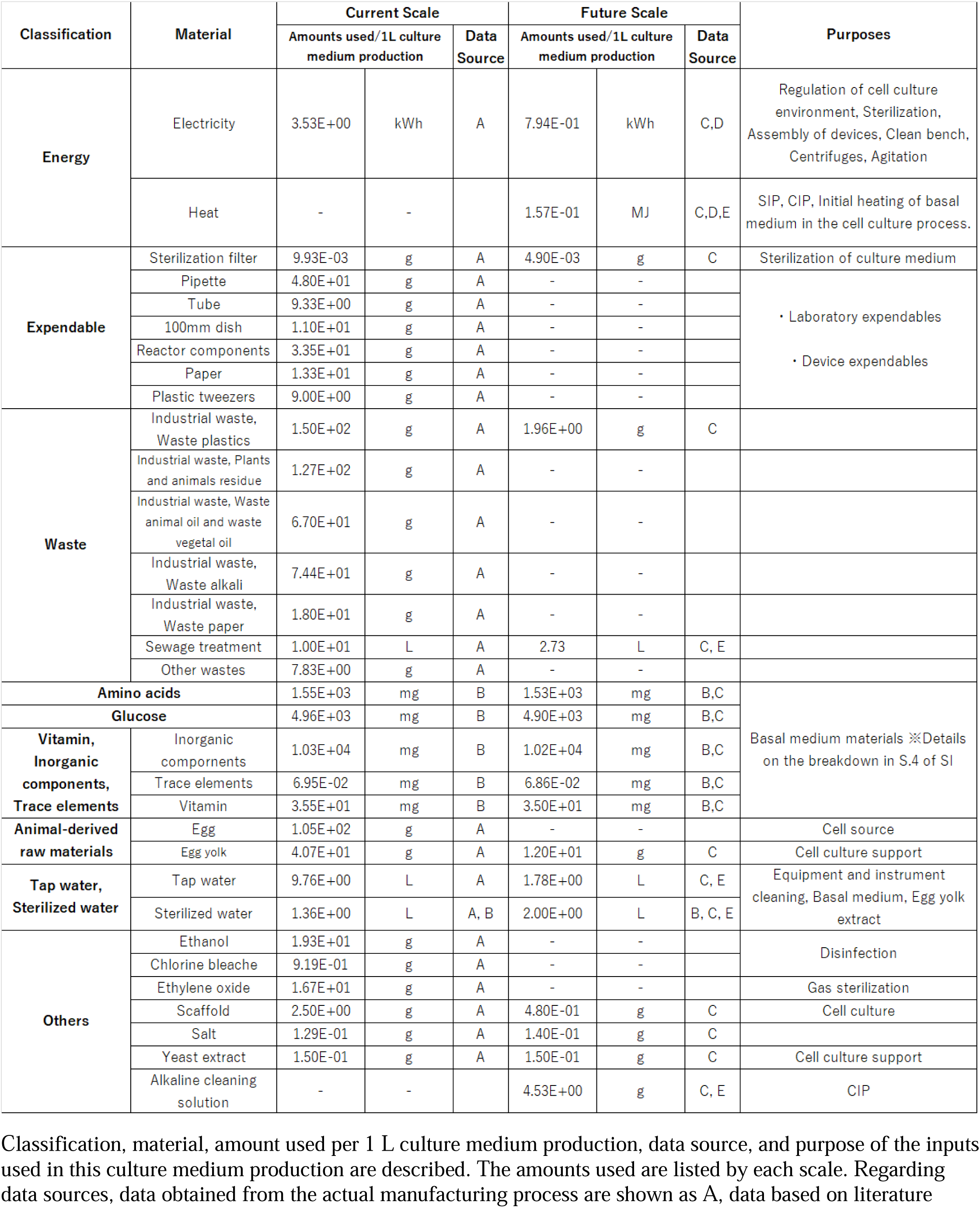
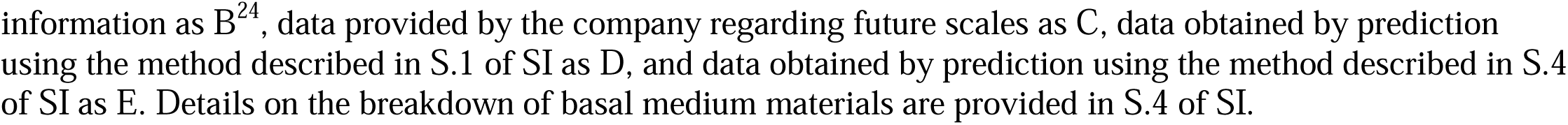
Inventory data for 1 L of the culture medium production.

| Classification | Material | Current Scale |  |  | Future Scale |  |  | Purposes |
| --- | --- | --- | --- | --- | --- | --- | --- | --- |
|  |  | Amounts used/1L culture medium production |  | Data Source | Amounts used/1L culture medium production |  | Data Source |  |
| Energy | Electricity | 3.53E+00 | kWh | A | 7.94E-01 | kWh | C,D | Regulation of cell culture environment, Sterilization, Assembly of devices, Clean bench, Centrifuges, Agitation |
|  | Heat | - | - |  | 1.57E-01 | MJ | C,D,E | SIP, CIP, Initial heating of basal medium in the cell culture process. |
| Expendable | Sterilization filter | 9.93E-03 | g | A | 4.90E-03 | g | C | Sterilization of culture medium |
|  | Pipette | 4.80E+01 | g | A | - | - |  | • Laboratory expendables<br>• Device expendables |
|  | Tube | 9.33E+00 | g | A | - | - |  |  |
|  | 100mm dish | 1.10E+01 | g | A | - | - |  |  |
|  | Reactor components | 3.35E+01 | g | A | - | - |  |  |
|  | Paper | 1.33E+01 | g | A | - | - |  |  |
|  | Plastic tweezers | 9.00E+00 | g | A | - | - |  |  |
| Waste | Industrial waste, Waste plastics | 1.50E+02 | g | A | 1.96E+00 | g | C |  |
|  | Industrial waste, Plants and animals residue | 1.27E+02 | g | A | - | - |  |  |
|  | Industrial waste, Waste animal oil and waste vegetal oil | 6.70E+01 | g | A | - | - |  |  |
|  | Industrial waste, Waste alkali | 7.44E+01 | g | A | - | - |  |  |
|  | Industrial waste, Waste paper | 1.80E+01 | g | A | - | - |  |  |
|  | Sewage treatment | 1.00E+01 | L | A | 2.73 | L | C, E |  |
|  | Other wastes | 7.83E+00 | g | A | - | - |  |  |
| Amino acids |  | 1.55E+03 | mg | B | 1.53E+03 | mg | B,C | Basal medium materials ※Details on the breakdown in S.4 of SI |
| Glucose |  | 4.96E+03 | mg | B | 4.90E+03 | mg | B,C |  |
| Vitamin, Inorganic components, Trace elements | Inorganic components | 1.03E+04 | mg | B | 1.02E+04 | mg | B,C |  |
|  | Trace elements | 6.95E-02 | mg | B | 6.86E-02 | mg | B,C |  |
|  | Vitamin | 3.55E+01 | mg | B | 3.50E+01 | mg | B,C |  |
| Animal-derived raw materials | Egg | 1.05E+02 | g | A | - | - |  | Cell source |
|  | Egg yolk | 4.07E+01 | g | A | 1.20E+01 | g | C | Cell culture support |
| Tap water, Sterilized water | Tap water | 9.76E+00 | L | A | 1.78E+00 | L | C, E | Equipment and instrument cleaning, Basal medium, Egg yolk extract |
|  | Sterilized water | 1.36E+00 | L | A, B | 2.00E+00 | L | B, C, E |  |
| Others | Ethanol | 1.93E+01 | g | A | - | - |  | Disinfection |
|  | Chlorine bleache | 9.19E-01 | g | A | - | - |  |  |
|  | Ethylene oxide | 1.67E+01 | g | A | - | - |  |  |
|  | Scaffold | 2.50E+00 | g | A | 4.80E-01 | g | C | Cell culture |
|  | Salt | 1.29E-01 | g | A | 1.40E-01 | g | C |  |
|  | Yeast extract | 1.50E-01 | g | A | 1.50E-01 | g | C | Cell culture support |
|  | Alkaline cleaning solution | - | - |  | 4.53E+00 | g | C, E | CIP |

Classification, material, amount used per 1 L culture medium production, data source, and purpose of the inputs used in this culture medium production are described. The amounts used are listed by each scale. Regarding data sources, data obtained from the actual manufacturing process are shown as A, data based on literature
information as B<sup>24</sup>, data provided by the company regarding future scales as C, data obtained by prediction using the method described in S.1 of SI as D, and data obtained by prediction using the method described in S.4 of SI as E. Details on the breakdown of basal medium materials are provided in S.4 of SI.

We used the LCI database IDEA Ver. 3.3 (IDEA) for the LCI database.^30^ IDEA is the most comprehensive inventory database available in Japan. IDEA shows the environmental burden intensity for each product, which traces back to the upstream processes, specifically, “from cradle to gate.” In this study, the LCI quantified the environmental load substances generated in this culture medium production by multiplying the amount of each material used and the waste generated, as presented in Table 4, by the corresponding environmental burden intensity in IDEA. “Current Energy Mix” used in this study’s [Current Scale] and [Future Scale, 2020 mix] was based on the “electricity, Japan, FY2020” environmental burden intensity because it was the latest data in IDEA.

IDEA provides information on the environmental burden intensity for generating 1 kWh of electricity using each electricity generation source. By multiplying these by the energy source composition proportion in Table 2, we calculated the environmental burden intensity for the “Electricity Mix in 2030” and “Renewable Electricity (RE) Mix.” Detailed life cycle inventory data were shown in S.5 of the SI.

### 2.6 Life Cycle Impact Assessment (LCIA)

In this study, the Life-cycle Impact assessment Method based on Endpoint modeling 2 (LIME2), an LCIA methodology that reflects the environmental conditions of Japan, was used in the LCIA.^31^ The environmental impact is quantified by multiplying the amount of each environmental burden calculated in the LCI by the LCIA coefficient and classifying the amount of each environmental burden with the impact category. This study performed LCIA for the following eight impact categories considered particularly important for culture medium production: climate change, ozone layer depletion, acidification, eutrophication, land use (occupation), land use (transformation), resource consumption, and water resources consumption. The results for all impact categories evaluated in LIME2 are shown in S. 6 of the SI.

## 3. RESULTS

Figure 3 shows the environmental impact of each material at current and future scales. Table 5 shows the environ

**Figure 3.**
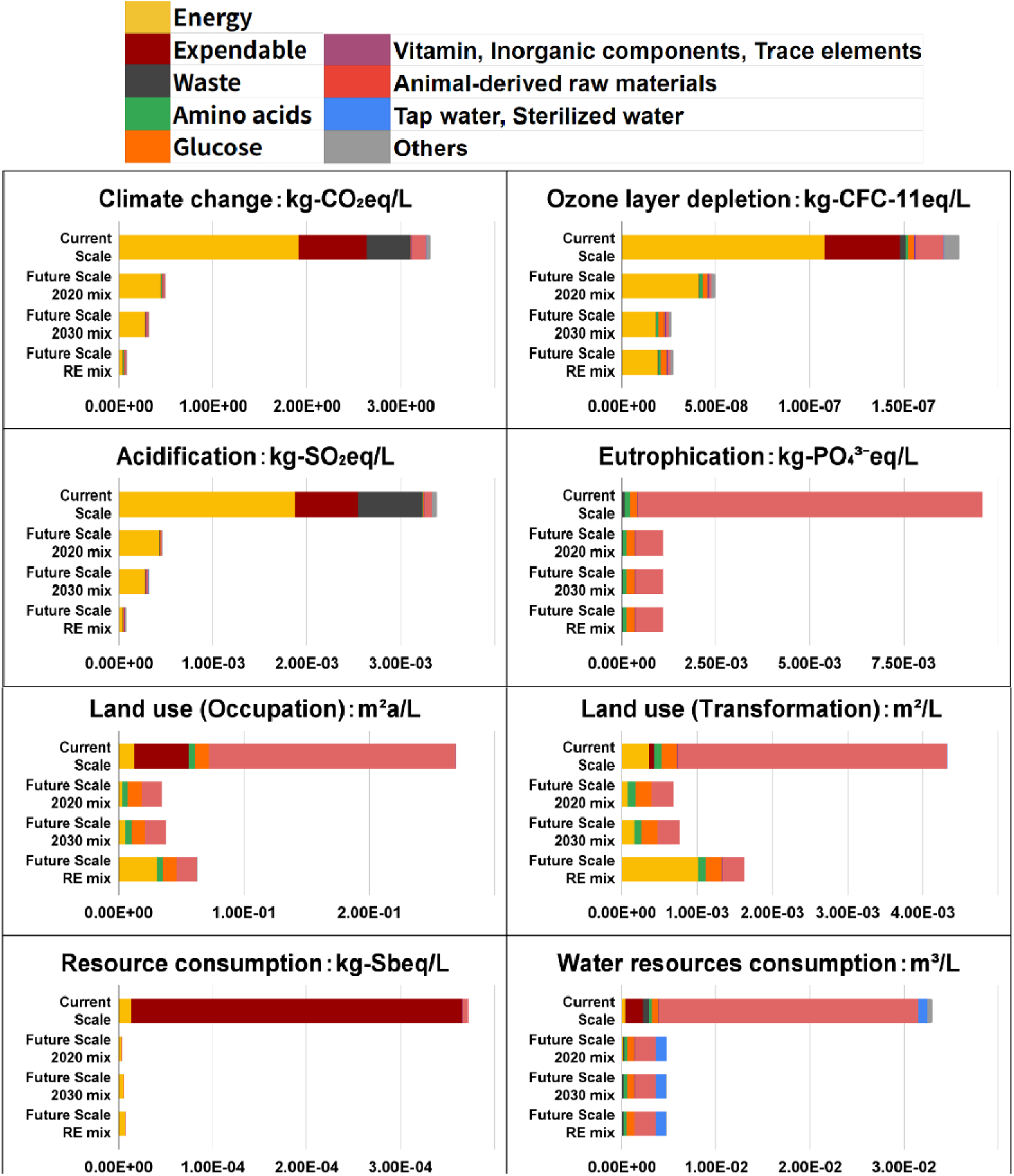
Environmental impact of each material for 1 L culture medium production.

**Table 5.** Environmental impact of 1 L culture medium production.

| Impact Category | Unit | Current Scale | Future Scale, 2020 mix | Future Scale, 2030 mix | Future Scale, RE mix |
| --- | --- | --- | --- | --- | --- |
| Climate change | kg-CO<sub>2</sub>eq | 3.32E+00 | 4.93E-01 | 3.20E-01 | 8.72E-02 |
| Ozone layer depletion | kg-CFC-11eq | 1.80E-07 | 4.96E-08 | 2.64E-08 | 2.75E-08 |
| Acidification | kg-SO<sub>2</sub>eq | 3.38E-03 | 4.66E-04 | 3.16E-04 | 7.73E-05 |
| Eutrophication | kg-PO<sub>4</sub><sup>3-</sup>eq | 9.61E-03 | 1.11E-03 | 1.11E-03 | 1.11E-03 |
| Land use (Occupation) | m<sup>2</sup>a | 2.69E-01 | 3.49E-02 | 3.76E-02 | 6.29E-02 |
| **Land use**<br>**(Transformation)** | m<sup>2</sup> | 4.34E-03 | 6.86E-04 | 7.73E-04 | 1.63E-03 |
| **Resource**<br>**consumption** | kg-Sbeq | 3.72E-04 | 3.89E-06 | 6.20E-06 | 7.89E-06 |
| **Water Resources**<br>**Consumption** | m<sup>3</sup> | 3.30E-02 | 4.74E-03 | 4.72E-03 | 4.71E-03 |

### 3.1 Current Scale

Energy accounts for over 50% of climate change, ozone layer depletion, and acidification. Animal-derived raw materials accounted for 73.1-95.6% of eutrophication, land use (Occupation and Transformation), and water resources consumption. Expendables accounted for a large proportion (16-94.7%) in the six categories except land use (transformation) and eutrophication. Therefore, electricity, animal-derived raw materials, and expendables are hotspots in the environmental impact of the current culture-medium production.

### 3.2 Future Scale

[Future Scale, 2020 mix] reduced environmental impacts by more than 70% in all impact categories compared to [Current Scale]. Among the categories, resource consumption had the largest rate of decline (99%), and ozone layer depletion had the smallest rate of decline (72.4%). [Future Scale, 2020 mix] results showed that energy accounts for 79.1-92% of the impacts of climate change, ozone layer depletion, acidification, and resource consumption. It was assumed that animal-derived raw materials would decrease at a future scale.

However, they still account for a large proportion (43-68.4%) of eutrophication, land use (Occupation and Transformation), and water resources consumption. Amino acids accounted for 10.6-13.9% of eutrophication and land use (Occupation and Transformation), and glucose accounted for 15.4-30.7% of eutrophication, land use (Occupation and Transformation), and water resources consumption. Therefore, energy, animal-derived raw materials, amino acids and glucose are hotspots for the environmental impact of the future scale.

[Future Scale, 2030 mix] and [Future Scale, RE mix] showed little change in eutrophication and water resources consumption, compared with [Future Scale, 2020 mix]. Climate change and acidification reduced by more than 30% in [Future Scale, 2030 mix] and by more than 80% in [Future Scale, RE mix], compared with [Future Scale, 2020 mix]. Ozone layer depletion is reduced by approximately 45% in both scenarios. In resource consumption and land use (Occupation and Transformation), [Future Scale, 2030 mix] and [Future Scale, RE mix] had larger impacts than [Future Scale, 2020 mix]. Resource consumption increased by 59.8% for [Future Scale, 2030 mix] and 103.5% for [Future Scale, RE mix]. Land use increased by 80.2% (Occupation) and 137.5% (Transformation) in [Future Scale, RE mix], while there was a slight increase in [Future Scale, 2030 mix].

## 4. Discussion

### 4.1 Current Scale

Electricity, animal-derived raw materials, and expendables are hotspots in the environmental impact of the current culture medium production.

Regarding the breakdown of electricity consumption, 30% was consumed by adjusting the culture environment, such as temperature and air control, in the cell culture process to prepare serum substitutes, and 28% was consumed by sterilizing instruments and equipment. Thus, reducing electricity consumption through these processes is important. Cell culture is common in cultivated meat production, and considerable studies are being conducted to improve cell culture efficiency.^32–34^ As the cell culture efficiency increases, the electricity consumption of this process is expected to decrease. In addition, the benefits of scale can be realized by scaling up, such as future scale; this is expected to reduce the electricity required for the cell culture process per liter of culture medium production. Besides, because this process is performed on a laboratory scale, the equipment and experimental apparatus are divided into smaller pieces, just as a bioreactor or other device comprises several parts. They must be sterilized individually, which tends to increase the number of sterilization procedures required. As technology becomes more established, these parts will be integrated into the device, and the number of sterilizations will be reduced. In sterilization, the larger the weight of the sterilization target, the lower the electricity consumption per unit weight.^35^ Therefore, the energy required for sterilization should be reduced by establishing the technology and decreasing the number of sterilization cycles.

Animal-derived raw materials were indicated as hotspots, particularly in eutrophication, land use (Occupation and Transformation), and water resources consumption. Eggs as a cell extraction source accounted for 72.1% of the animal-derived raw materials used in the current scale. Therefore, the cell extraction source is required to be changed. Using cell lines is one of the methods to obtain cells without slaughtering the animals. As cell lines divide semi-permanently, it is possible to easily prepare cells in large quantities, which is expected to significantly reduce the use of animal-derived raw materials.^26^ In addition, animal welfare is generally recognized as one of the advantages of cultivated meat. It is necessary to avoid using animal-derived raw materials in the production of culture medium from the perspective of environmental impact shown in this study and animal welfare and consumer acceptance.^36^ Therefore, there is a need to transition to using cell lines as a cell source or collecting biopsy cells from low-environmental-impact livestock without slaughter.^37,38^ Expendables accounted for a large proportion of the seven impact categories except for eutrophication, and expendables increased the amount of waste. Therefore, it is essential to reduce the use of expendable products. Expendables include equipment such as conductors for connecting multiple bioreactors and experimental instruments such as pipettes and tubes. Equipment expendables are expected to be reduced by integrating all equipment parts and making them multi-usable. In addition, expendables should be reduced through mechanization of the process, and the amount of expendables used per 1L of culture medium production should be reduced by improving the productivity of one batch.

### 4.2 Future Scale

Energy, animal-derived raw materials, amino acids and glucose were assumed to be hotspots in the environmental impact of future culture medium production.

The use of animal-derived raw materials was assumed to decrease because of a change in the cell extraction source from eggs to cell lines and productivity improvements. However, egg yolk extract for assistance in the cell culture process will be used on a future scale, and animal-derived raw materials were assumed to account for a large proportion of eutrophication, land use (Occupation and Transformation), resource consumption, and water resources consumption. Further, as mentioned above, animal-derived raw materials should be avoided from the perspective of environmental impact, animal welfare, and consumer acceptance. In the cell culture process for preparing serum substitutes, further research and development are needed in a direction that does not require animal-derived raw materials. One specific strategy is to use a serum substitute component prepared in a different batch as a culture aid instead of egg yolk extract.

It is also assumed that with the transition from the current to the future scale, expendables and waste will decrease because of the multi-use of the entire equipment and increased production of culture medium per batch. The reduction in the use of expendable, waste, and animal-derived raw materials has led to a larger contribution of amino acids and glucose, the raw materials of target basal medium in the eutrophication, water resources consumption, and land use. This is due to the water and land use and run-off of fertilizer-derived nitrogen and phosphorus from the soil to the hydrosphere during the production of grains, which are the main extraction source of amino acids and glucose.^39–41^ Therefore, a shift to algae and other extraction sources with lower environmental impact is necessary to further reduce the environmental impact of culture medium production in the future.^42^

Energy is assumed to account for a large share of the impact categories. Therefore, scenarios [Future Scale, 2030 mix] and [Future Scale, RE mix] were considered with different electricity sources at future scales. For the main changes in the electricity source composition, in [Future Scale, 2030 mix], it is assumed that the share of solar and nuclear power will increase and that the share of coal and Liquefied Natural Gas (LNG) will decrease compared to [Future Scale, 2020 mix]. The electricity composition of [Future Scale, RE mix] is assumed to consist only of solar and wind power, eliminating coal, LNG, nuclear power, hydroelectric power, and other sources accounting for a large share of the electricity composition in the other two scenarios. The share of renewable energy in electricity composition is assumed to increase in ascending order: [Future Scale, 2020 mix], [Future Scale, 2030 mix], and [Future Scale, RE mix]. As a result of these changes, the impact of ozone layer depletion decreased in [Future Scale, 2030 mix] and [Future Scale, RE mix], compared with [Future Scale, 2020 mix], but increased slightly from [Future Scale, 2030 mix] to [Future Scale, RE mix]. Resource consumption increased in both scenarios, compared with [Future Scale, 2020 mix]. However, their increase is small compared to the environmental impact of each category of the current scale. Impacts on climate change and acidification decreased considerably as the share of renewable energy in electricity composition increased.

However, land use, both occupation and transformation, notably increased as the share of renewable energy increased. In particular, focusing only on the impact of energy on land use, there was an increase of more than 1,000% in occupation and transformation from [Future Scale, 2020 mix] to [Future Scale, RE mix].

The transition to renewable energy is essential for reducing the environmental impact of this culture medium, sustainable development, and solving various current environmental problems.^43^ However, the results of this study indicated that an increase in the percentage of renewable energy in the electricity source mix used in the culture medium production considerably increases the impact on land use. The impact of renewable energy on land use has been discussed in many previous studies.^44–46^ Increased land use may lead to greenhouse gas emissions due to the loss of primary plant production and loss of biodiversity due to habitat loss, change, and fragmentation.^47^ It may also undermine the advantage of cultivated meat, which requires less land use for meat production compared to livestock production, as shown in previous studies.^6–8^ One way to address such issues related to renewable energy and land use is to use wastelands, building roofs, urban areas, and water surfaces as sites for power generation facilities or share the land with other land uses such as agricultural land and pasture land.^46–48^ Among these measures, those that are not difficult to adapt to culture medium production may include installing electricity generation facilities on the rooftops of production plants and sharing land by installing them within production sites. However, there is potential for debate regarding land use for renewable energy, as some argue that renewable energy uses less land than other forms of energy because the efficiency of power generation varies depending on local environmental characteristics, such as wind and solar radiation, and because additional fuel extraction and transportation are not required.^49,50^ The priority is reducing electricity consumption to reduce the environmental impact of culture medium production. The results of this study indicate that the cell culture process for the preparation of serum substitutes consumes the most electricity at current and future scales. As described above, as the efficiency of cell culture improves, electricity consumption in this process and the overall production of the culture medium are also expected to be reduced. Therefore, it is possible that the benefits of scale can be realized if the scale is further increased from that assumed in this study. Thus, it is important to recognize that while using renewable energy in production is essential for producing this culture medium, it also involves issues such as land use. The first goal is to reduce electricity consumption. Comprehensively considering various environmental impacts and methods of installing power generation facilities is also necessary.

### 4.3 Basal Medium and Serum Substitute Components

Figure 4 shows the environmental impact of each material in the production of 0.5 L basal medium (BM) and 0.5 L serum substitute components (SS) in all scenarios, because this culture medium 1L is prepared by mixing 0.5 L of serum substitute components and 0.5 L of basal medium. Basal medium is used to produce the serum substitute components of this culture medium. To avoid confusion, the environmental impact of the basal medium in the production of serum substitute components is shown as the basal medium alone, not broken down into inputs. (The results for basal medium 0.5 L production in Figure 4 do not include the basal medium used to produce the serum substitute component.)

**Figure 4.**
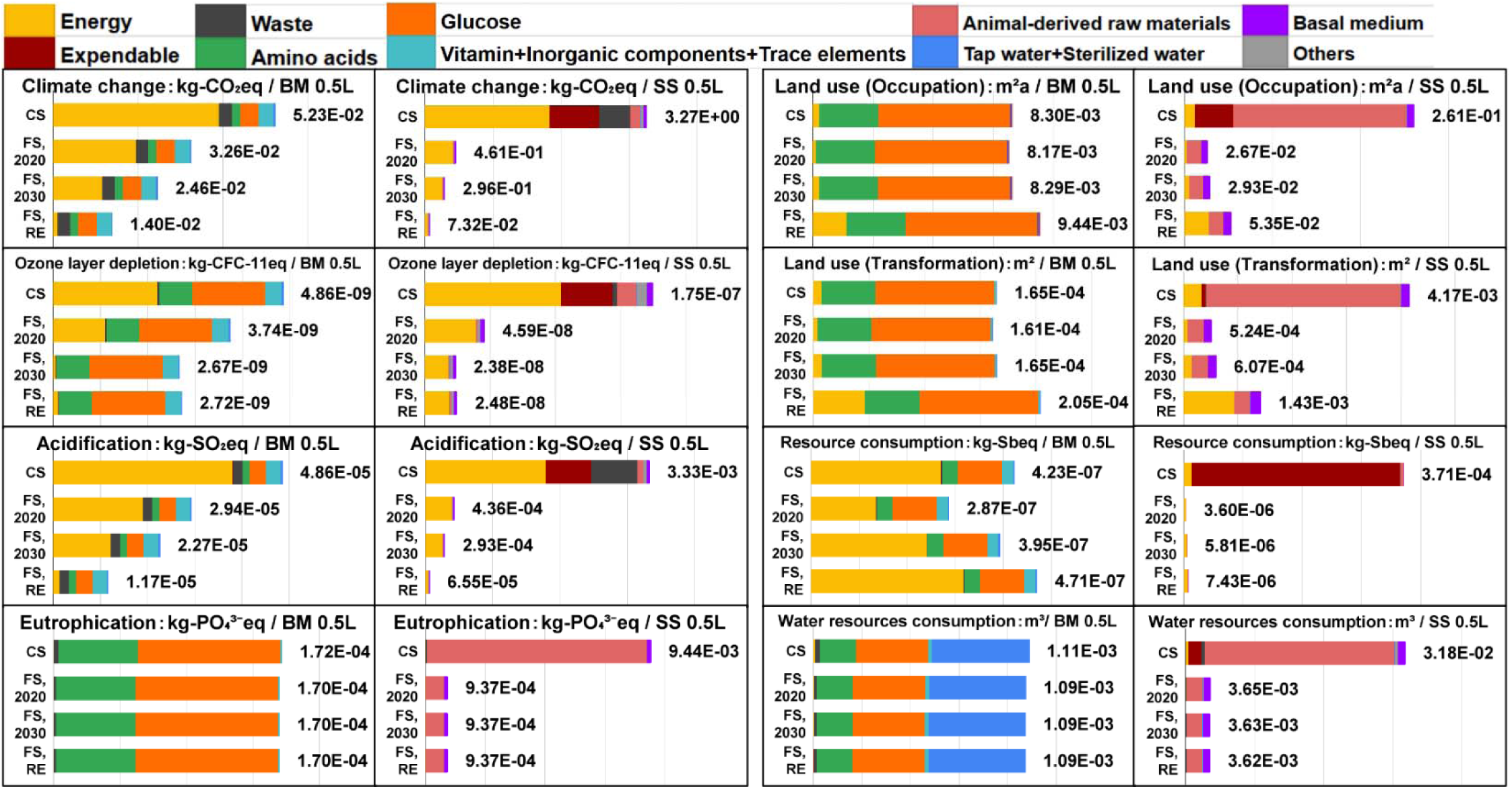
Environmental impact of each material for 0.5 L basal medium (BM) production and serum substitute components (SS) production. This study’s culture medium 1 L is prepared by mixing 0.5 L of basal medium and 0.5 L of serum s bstitute component. BM: basal medium, SS: serum substitute components, CS: Current Scale, FS, 2020: Future Scale 2020 mix, FS, 2030: Future Scale 2030 mix, FS, RE: Future Scale RE mix.

In all categories, serum substitute components accounted for more than 96.2% (at the current scale) and more than 76.6% (at the future scale) of the environmental impact of culture medium production.

In the production of basal medium, amino acids and glucose accounted for a large proportion of land use (Occupation and Transformation) and eutrophication. Tap water and sterilized water accounted for more than 45% of water resources consumption, as it was used to produce basal medium. Energy was mainly used during the sterilization of bottles for stocking basal medium and filter sterilization. Quantities of amino acids and glucose were larger than those of vitamins. The amount of inorganic salts was larger than those of amino acids and glucose. Yet the environmental impact of grain production as a raw material of amino acids and glucose was larger. Therefore, the hotspots of the environmental impact of basal medium production were indicated to be energy, amino acids, and glucose.

The environmental impact of each material in serum substitute components production showed a similar trend to that of the target culture medium. Energy, animal-derived raw materials, and expendables accounted for a large proportion of the environmental impact of serum substitute components production at the current scale. In contrast, energy and animal-derived raw materials had a large contribution at the future scale.

### 4.4 Comparison of Previous Studies

Table 6 shows a summary of previous LCA studies of culture medium production. Nikkhah et al. 2024 ^21^ performed LCA of Beefy-9 and Beefy-R production, animal component-free culture media using DMEM/F-12 as basal medium, recombinant growth factors and recombinant proteins as serum substitute components. Wali et al. 2024 ^11^ performed LCA of culture medium containing FBS and alternative FBS-free media containing serum substitutes, protein hydrolysates, and recombinant growth factors. As the LCIA methods used in this study and previous studies differ, the methods used to assess each impact category differ. Thus, results were not simply compared quantitatively. However, regarding the impact category of climate change/global warming potential, both LCIA methods are based on the Intergovernmental Panel on Climate Change (IPCC) Fifth Assessment Report’s 100-year time horizon global warming potentials (GWP).^31,51,52^ Therefore, only the climate change/GWP results were compared quantitatively.

**Table 6.** Summary of previous LCA studies of culture medium production.

| Authors (year) |  | This study |  |  |  | Nikkhah et al. (2024)<sup>21</sup> |  | Wali et al. (2024)<sup>11</sup> |  |  |  |
| --- | --- | --- | --- | --- | --- | --- | --- | --- | --- | --- | --- |
| System boundary |  | Cradle-to-gate |  |  |  | Cradle-to-factory gate |  | Cradle-to-gate |  |  |  |
| Functional unit |  | 1L of culture medium |  |  |  | 1L of culture medium |  | 1L of culture medium |  |  |  |
| LCI database |  | LCI database IDEA Ver. 3.3 |  |  |  | Ecoinvent 3 and Agri-footprint 6 |  | Ecoinvent 3.8 and Agri-footprint |  |  |  |
| LCIA methodology |  | Life-cycle Impact assessment Method based on Endpoint modeling 2 |  |  |  | ReCiPe 2016 Midpoint (H) |  | ReCiPe 2016 Midpoint (H) and cumulative energy demand |  |  |  |
| Scenario / Electricity grid (Sensitivity analysis) |  | Current Scale | Future Scale, 2020 mix | Future Scale, 2030 mix | Future Scale, RE mix | Global average electricity grid (electricity grids of the United States, Singapore, and Europe) |  | Electricity grids of Finland (Denmark, Norway and renewable energy) |  |  |  |
| Basal medium |  | Food ingredients basal medium |  |  |  | DMEM/F-12 |  | DMEM/F-12 |  |  |  |
| Serum substitution method |  | By co-culturing multiple organ cells the interaction of each organ cell secretes serum substitute components. |  |  |  | Recombinant growth factors and proteins (Mainly recombinant albumin) | Recombinant growth factors and proteins (Mainly rapeseed protein isolates) | 10% FBS | 2% growth factors + 2% FBS | 2% Essential 8™ + protein hydrolysates | Tri-basal 2.0 + ITS |
| Climate change / Global warming potential (GWP) |  | 3.32 kgCO<sub>2</sub>eq | 4.93× 10<sup>-1</sup> kgCO<sub>2</sub>eq | 3.2× 10<sup>-1</sup> kgCO<sub>2</sub>eq | 8.72× 10<sup>-2</sup> kgCO<sub>2</sub>eq | 3.90× 10<sup>-1</sup> kgCO<sub>2</sub>eq | 8.22× 10<sup>-2</sup> kgCO<sub>2</sub>eq | 1.33× 10<sup>-1</sup> kgCO<sub>2</sub>eq | 4.61× 10<sup>-2</sup> kgCO<sub>2</sub>eq | (2.47-3.47)× 10<sup>-2</sup> kgCO<sub>2</sub>eq | 2.93× 10<sup>-2</sup> kgCO<sub>2</sub>eq |
| Main contributor to each impact | GWP | Energy |  |  |  | Recombinant proteins and growth factors |  | FBS | FBS | Amino acids | Amino acids |
|  | Water consumption | Animal-derived raw materials |  |  |  | Recombinant proteins and growth factors | Basal medium | Water for basal medium |  |  |  |
|  | Land use | Animal-derived raw materials | Animal-derived raw materials, Glucose | Animal-derived raw materials, Glucose | Energy | Recombinant proteins and growth factors, Basal medium | Basal medium | Glucose | Glucose, Amino acids | Glucose, Amino acids | Amino acids |
|  | (Terrestrial) acidification | Energy |  |  |  | Recombinant proteins and growth factors | Basal medium | FBS | FBS | Raw material, Amino acids | Amino acids |
|  | Freshwater eutrophication | Animal-derived raw materials |  |  |  | Recombinant proteins and growth factors | Basal medium | FBS | Growth factors | Raw material, Amino acids, Chemicals & other components | Amino acids |
|  | Marine eutrophication |  |  |  |  | Basal medium |  | FBS | FBS | Raw material, Amino acids, Glucose | Growth factors |

The large contribution of the environmental impact of amino acid and glucose production among the basal medium raw materials was common to all the studies. It is also common for basal medium production to account for a large proportion of land use, eutrophication, and water resources consumption. The trend towards increased land use due to renewable energy was also common with Nikkhah et al. 2024 ^21^. Compared with previous studies, the proportion of animal-derived raw materials in the environmental impact of this study’s culture medium production was remarkable.

Regarding climate change/GWP, this study’s [Current Scale] results were overwhelmingly higher than those of previous studies, and [Future Scale, 2020 mix] results were also higher. [Future Scale, 2030 mix] result had a slightly lower impact than Beefy-9 (Nikkhah et al. 2024 ^21^) and [Future Scale, RE mix] result was lower than DMEM/F12+10% FBS (Wali et al. 2024 ^11^), and closer to Beefy-R (Nikkhah et al. 2024 ^21^). (The results of Wali et al. 2024 ^11^ had a smaller impact among all studies partly because the Finnish electricity grid’s environmental impact is lower than the others.) One of the reasons for this is that, in all impact categories, not just climate change, this study’s culture medium accounted for a larger proportion of serum substitute components compared to previous studies; this is because in the culture medium of the previous study, the basal medium accounted for more than 90% of 1 L of culture medium, to which the serum substitute components were added. In contrast, in this study’s culture medium, the serum substitute components accounted for 50% of the culture medium. In addition, a major feature of this culture medium is that the production process includes a cell culture process, and this process accounts for a large proportion of the environmental impact of this culture medium production. As described above, the improved efficiency of cell culture and further scale-up in this process is expected to further reduce the environmental impact. The energy used in this cell culture process accounts for a large proportion of the environmental impact of this culture medium production. Therefore, the environmental impact of this culture medium depends more on the environmental impact of the energy source than other culture mediums. As the [Future Scale, 2030 mix] has a relatively large impact among the previous study results, it is not sufficient to follow the changes in the policy target of Japanese electricity grid. Therefore, the importance of the energy source used in the production process has been mentioned in previous studies, particularly in this study’s culture medium production, it is necessary to use a sustainable electricity grid through independent efforts.

### 4.5 Sensitivity Analysis

This study’s culture medium contained 15 amino acids. Yet data on the environmental burden intensity of individual amino acids could not be obtained from the IDEA used as an LCI database. Therefore, data on the environmental burden intensity of “edible amino acids” were used to assess all amino acids (Details in S.4 of SI). However, the environmental impact of production varies with the kind of amino acid. Amino acids are listed as a hotspot in the future scale of culture medium production in this study, and uncertainty in the environmental impact of amino acid production may affect the overall environmental impact of culture medium production. IDEA has data on “ monosodium glutamate” besides “edible amino acids” as amino acid production. Therefore, a sensitivity analysis was conducted in this study by examining the impact on the environmental impact of culture medium production when using data on “monosodium glutamate” instead of “edible amino acids” to assess amino acid production.

Figure 5 shows the difference in the environmental impact of culture-medium production in each scenario, using data on “monosodium glutamate” instead of “edible amino acids” to assess amino acid production. The sensitivity analysis showed that impact categories of climate change, ozone layer depletion, acidification, and resource consumption were largely affected by changes in the environmental burden intensity of amino acid production. The most highly affected scenarios differed depending on the impact category. [Future Scale, RE mix] had the highest impact on climate change and acidification, with 4.1 and 6.3% increases, respectively. [Future Scale, 2030 mix] had the largest impact on ozone layer depletion, with an increase of 2.7%. Regarding resource consumption, [Future Scale, 2020 mix] had the highest impact, with 2.8% increase. Conversely, eutrophication, land use, and water resources consumption categories were small affected by changes in the environmental burden intensity of amino acid production.

**Figure 5.**
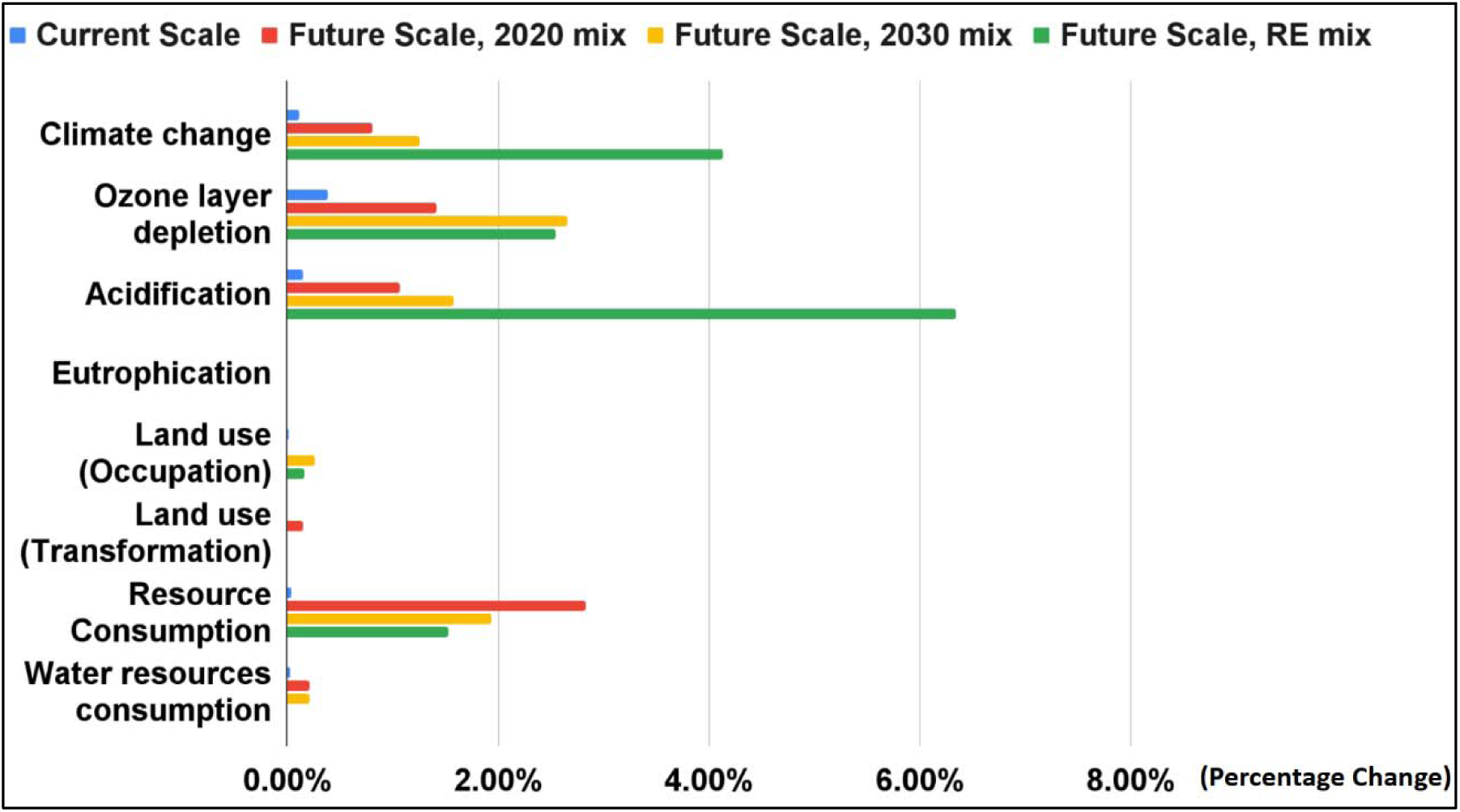
Sensitivity analysis results: The percentage change in the environmental impact of culture medium production for difference in the environmental burden intensity of amino acid production.

However, no change was observed in the magnitude relationship between scenarios owing to a change in the environmental burden intensity of amino acid production. Therefore, the results of this study were relatively stable, with changes in the environmental burden intensity of amino acid production. Meanwhile, as mentioned in previous studies, the accumulation of detailed environmental impact knowledge of amino acid and vitamin production is needed for accurate LCA study of culture media and cultivated meat production in the future.^11,21^

### 4.6 Limitations and Future Prospects of This Study

This study assessed the environmental impact of the company’s production of culture medium consisting of serum-free, food and complex ingredients at current and future scales. Consequently, the environmental impact of culture medium production was expected to decrease considerably with the transition from current to future scale. The hotspots for environmental impacts of culture medium production were electricity, animal-derived raw materials, and expendables at the current scale, and energy, animal-derived raw materials, amino acids, and glucose at the future scale.

The data used in this study were collected from only one company because there is currently only one xample of a culture medium consisting of serum-free, food and complex ingredients produced using cell interaction. Thus, the results of this study may differ from those of LCA for other cases using the same culture medium type. Therefore, LCAs should be conducted if other cases related to the same culture medium type arise. Moreover, although this study conducted an LCA only on the culture medium production process, the ultimate purpose of the culture medium under study was to cultivate meat. An LCA covering the entire cultivated meat production process will be necessary in the future, requiring the findings of culture medium production obtained in the present study.

In addition, this study’s culture medium contained animal-derived raw materials at current and future scales. However, from the perspective of environmental impact, animal welfare, and consumer acceptance, animal-derived raw materials should be avoided in culture medium production for cultivated meat. Animal welfare is one of the motivations for cultivated meat, and it is necessary to establish evaluation methods and implement evaluations that include animal welfare. The production cost is also an important factor for the industrialization of cultivated meat, and cost aspects need to be assessed through techno-economic analysis. Therefore, a comprehensive assessment is needed that includes environmental impacts and social, cost, and animal welfare aspects.

## ASSOCIATED CONTENT

### Supporting Information

S.1 Cell culture by this study’s culture medium and other culture media.; S.2. Prediction of energy consumption of bioreactors used at future scales.; S.3. Prediction of energy and input consumption of cleaning-in-place (CIP) and steam-in-place (SIP).; S.4. Details of the raw materials of the basal culture medium.; S.5. Detailed life cycle inventory data.; S.6. Life Cycle Impact Assessment (LCIA).

## AUTHOR INFORMATION

### Author Contributions

Conceptualization: [Natsufumi Takenaka, Kimiko Hong-Mitsui, Kazuhiro Kunimasa, Kotaro Kawajiri, Chihiro Kayo, Naoki Yoshikawa]; Methodology: [Natsufumi Takenaka, Kotaro Kawajiri, Chihiro Kayo, Naoki Yoshikawa]; Investigation: [Natsufumi Takenaka, Kimiko Hong-Mitsui, Kazuhiro Kunimasa]; Writing – original draft preparation: [Natsufumi Takenaka, Chihiro Kayo, Naoki Yoshikawa]; Writing – review and editing: [Kimiko Hong-Mitsui, Kazuhiro Kunimasa, Kotaro Kawajiri]

## ACKNOWLEDGMENTS

The authors thank the Tokyo University of Agriculture and Technology for their generous support. The authors are also grateful to IntegriCulture Inc. for providing the data. This research was funded by IntegriCulture Inc. We would like to thank Editage (www.editage.jp) for English language editing and making the graphical abstract.

## ABBREVIATIONS

LCA, life cycle assessment; FBS, Fetal Bovine Serum; ISO, International Organization for Standardization; DMEM, Dulbecco’s Modified Eagle’s Medium; SI, Supporting Information; CIP, Cleaning in Place; SIP, Sterilization in Place; LNG, Liquefied Natural Gas; RE, Renewable Energy; LCI, Life Cycle Inventory; IDEA, LCI database IDEA Ver. 3.3; LCIA, Life Cycle Impact Assessment; LIME2, Life-cycle Impact assessment Method based on Endpoint modeling 2; BM, Basal medium; SS, Serum substitute components; CS, Current Scale; FS, 2020, Future Scale, 2020 mix; FS, 2030, Future Scale, 2030 mix; FS, RE, Future Scale, RE mix; IPCC, Intergovernmental Panel on Climate Change; GWP, global warming potential.

