## Supporting Information for "Environmental Impacts of Producing Culture Medium Consisting of Serum-free, Food and Complex Ingredients for Cultivated Meat"

Corresponding Author

Summary of Supporting Information Content:

S.1 Cell culture by this study's culture medium and other culture media.

S.2. Prediction of energy consumption of bioreactors used at future scales.

S.3. Prediction of energy and input consumption of cleaning-in-place (CIP) and steam-in-place (SIP).

S.4. Details of the raw materials of the basal culture medium.

S.5. Detailed life cycle inventory data

S.6. Life Cycle Impact Assessment (LCIA)

**S.1.** **Cell culture by this study's culture medium and other culture media.**

Table S.1 shows the cell viability of duck liver cells cultured for 2 days from the same initial cell number in this study’s basal medium only, this study’s basal medium/FBS 10%, and this study’s culture medium, with the number of cells surviving in this study’s basal medium/FBS 10% as 1 (n=3). The experiment measured the amount of ATP in the cells after 2 days of culture by CellTiter-Glo® 2.0 Cell Viability Assay kit (Promega, Madison, WI, USA).

Table S. 1. Results of cell culture for two days.

|  | Cell viability (ratio) |
| --- | --- |
| This study's basal medium | 0.0273592 |
| This study's basal medium / FBS 10% | 1 |
| This study's culture medium | 1.07249 |

**S.2.** **Prediction of energy consumption of bioreactors used at future scales.**

In the future scale, the cell culture process for the production of serum substitute components assumed the use of a 10,000 L adjustment reactor (Stirred-Tank Reactors (STR)) without cell culture and 100 L feeder reactors (STR)×3 with cell culture. The height, diameter, filling capacity and surface area of each reactor are listed in Table S. 2. The duration of one batch is 28 days (672 h). The temperature was assumed to be 16.2 ℃, the annual average temperature in Fujisawa, Kanagawa, Japan, where the laboratory is located.^1^ The temperature during culture was maintained at 37 ℃.

Table S. 2. Reactor geometry.

|  | **Adjustment reactor (10,000L)** | **Feeder reactor(100L)** | **Dara Source** |
| --- | --- | --- | --- |
| Height (m) | 3.7 | 0.8 | Mattick et al. 2015 ^2^, Tuomisto et al, 2011 ^3^ |
| Diameter (m) | 1.85 | 0.4 |  |
| Surface area (m^2^) | 26.9 | 1.26 |  |
| Filling capacity (%) | 50 | 50 | Data on equipment used by the company. |

Table S.2 lists the energy use of the reactors at the future scale. The methodology for estimating those energies is explained below.

Table S. 3. Energy consumption of reactors.

|  | **Electricity consumption for 4 weeks.** | **Heat consumption** | **Prediction method** |
| --- | --- | --- | --- |
| Feeder reactor (100 L) | 2,990 kWh | - | S.2.1^4,5^ |
| Feeder reactor (100 L) | 2,990 kWh | - | S.2.1^4,5^ |
| Feeder reactor (100 L) | 2,990 kWh | - | S.2.1^4,5^ |
| Adjustment reactor (10,000 L) | 7,321 kWh | - | S.2.2^6,7,8^ |
| Pump | 296 kWh | - | Data on equipment used by the company. |
| Initial heating | - | 522 MJ | S.2.3 |

**S.2.1 Electricity consumption of feeder reactor (100 L)**

The energy use sources of the feeder reactor were assumed to be heater, aeration, and mixing, and electricity was assumed to be the energy source. In this study, the energy consumption of the feeder reactor was estimated using the method of Kawajiri et al. 2020 ^4^ based on the electricity consumption data of the 15 L and 60 L STRs used in the target company. The electricity consumption of the reactors used for reference is listed in Table S. 4. By fitting the data in Table S.4 to the equation (1) given by Kawajiri et al. 2020 ^4^, the electricity consumption of feeder reactors for the heater and aeration was predicted. The scaling factor for heater and aeration was obtained by fitting the electricity consumption of heater and aeration of 15 L STR and 60 L STR to (1). Mixing power consumption quoted actual product data.^5^

Table S. 4. Electricity consumption of the reactors used for reference.

|  | **Heater** | **Aeration** | **Dara Source** |
| --- | --- | --- | --- |
| 15 L STR | 0.35 kW | 1.5 kW | Data on equipment used by the company. |
| 60 L STR | 1.0 kW | 2 kW |  |

**P=P_0_(S_X_/S_0_) ^f^** 　(1)

P： Electricity Consumption of Target Scale (100L feeder reactor)

P_0_： Electricity Consumption of Reference Scale (Reference scales in Table S.4)

S_X_： Volume of Target Scale (100L feeder reactor)

S_0_： Volume of Reference Scale (Reference scales in Table S.4)

f： Scaling factor (f _Heater_=0.76, f _Aeration_=0.21)

**S.2.2 Electricity consumption of adjustment reactor (10,000 L)**

The energy use sources for the adjustment reactor were assumed to be heater, aeration, and mixing, and electricity was assumed to be the energy source. Electricity consumption for the aeration and mixing of the adjustment reactor was based on data for a 10,000 L STR in Sinke. et al. 2023 ^6^.

On the contrary, as the adjustment reactor (10,000 L) in this study does not perform cell culture, the electricity consumption of the heater was estimated separately. Cell proliferation is an exothermic reaction. Therefore, in previous studies, the energy consumption of the heater of a bioreactor has been estimated based on the relation between heat loss to the surroundings and the heat of reaction for cell proliferation. ^6,7^　However, because the adjustment reactor (10,000 L) in this study does not perform cell culture, the energy to maintain 37 ℃ from the heat loss to the surroundings was estimated as the energy consumption of heat in the adjustment reactor (10,000 L). This maintain energy was assumed to come from electricity with a 100% energy conversion.^7^ The feeder reactor was assumed to regulate the reaction heat from cell proliferation. The heat loss to the surroundings was estimated in the following way according to the method of Tuomisto. et al. 2022 ^7^.

The heat loss to the surroundings, Q_L_ (W), was dependent on the surface area of the reactor vessel, A (m^2^), the vessel wall thickness, x (m) material (subscript: i) and consequent thermal conductivity of the material, k (W m^-1^ K^-1^) and the temperature difference between the reactor internals and the surrounding environment, ΔT (K), defined by Equation 2.^7^ The reactor vessels were assumed to be made of stainless steel (x_ss_ = 0.005 m and k_ss_ = 16 W m^-1^ K^-1^) with a layer of glass lagging for insulation (x_gw_ = 0.025 m and k_gw_ = 0.04 W m^-1^ K^-1^).^7,8^ The culture temperature is 37 ºC and the surrounding temperature is 16.2 ºC.

**Q_L_ = ΔT / Σ (x_i_ / A k_i_)　(2)**

**S.2.3 Energy consumption of initial heating of basal medium**

Initial heating of the culture medium during the preparation of serum substitute components was assumed to be the energy to warm 5000 L of basal medium from 16.2 ℃ to 37 ℃. The energy source was assumed to be heat energy. The thermal efficiency was assumed to be 83.5%.^9^

**S****.3.** **Prediction of energy and input consumption of cleaning-in-place (CIP) and steam-in-place (SIP).**

The energy used for CIP and SIP was assumed to be heat energy. The model did not reuse cleaning solutions and rinsing water. Table S.5 lists the energy and input use of the SIP and CIP in the Future scale.

**S.3.1 Prediction of energy and input consumption of CIP**

The inputs and energy used for CIP for reactor cleaning at the future scale were estimated according to the method of Tuomisto. et al. 2022 ^7^ using the following procedure.

The requirements for CIP have been evaluated using the methodology of the previous study.^2^ The cleaning stage was assumed using 1% w/v sodium hydroxide solution at 77.5 °C in a water volume equal to the total reactor volume plus 10% as a reserve for additional units. This is in line with recommendations for cleaning industrial bioreactors.^9^ In addition to the water required for the alkali cleaning solution, sterilized water is required for a 5- 6 minute pre-rinse and a post-clean rinse to remove residual alkali from the system.^10^ The energy required for CIP was assumed to be the energy required to increase the water temperature from 16.2 ℃ to 77.5 ℃ and calculated by equation (3). The thermal efficiency was assumed to be 83.5%.^9^

**E _CIP_ = m _water_ Cp_, water_ ∆T　(3)**

E _CIP_ (kJ)：The energy required for CIP

m _water_ (kg)：Mass of the 1% w/v sodium hydroxide solution 11,330L

Cp_, water_ (kJ kg^-1^ K^-1^)：The specific heat capacity of the 1% w/v sodium hydroxide solution

∆T (K)：the temperature difference between 16.2 ℃ and 77.5 ℃

**S.3.2 Prediction of energy and input consumption of SIP**

Inputs and energy to the SIP for sterilizing reactors were estimated with reference to Tuomisto. et al. 2022 ^7^ and Sinke. et al. 2023 ^6^. The SIP was assumed to be carried out under conditions of 121.5 ℃ for 20 min, using 100 kg of steam per cubic meter of working volume. Q _steam gen_ (kJ) as the energy required for SIP to generate the steam per batch was calculated using equation (4). Equation (5) was used to determine Q_SS, heat up_ (kJ) as the energy required to heat the stainless steel reactor vessels with m _Reactors_ (kg) as the mass of the reactor vessels was 1740 kg (inclusive of 10% design contingency for piping and instrumentation), C_P, SS_ (kJ kg^-1^ K^-1^ ) as the specific heat capacity of stainless steel taken as 0.45 kJ kg^-1^ K^-1^ and ∆T (K) as the temperature difference between 16.2 ℃ and 121.5 ℃, Q_SS, Losses_ (kJh) is the rate of heat loss to the surrounding (it is assumed by equation (2)), t _SIP_ (h) is the period for SIP assumed to be 20 min (0.33 h) and η_Steam generation_ (%) is the efficiency of steam generation, assumed to be 80% ^11^.

**Q _steam gen_ =［Q_SS, heat up_ + (Q_SS, Losses_ t _SIP_)］/ η_Steam generation_　(4)**

**Q_SS, heat up_ = m _Reactors_ C_P, SS_ ∆T　(5)**

Table S. 5. Energy and input consumption of CIP and SIP.

|  | **Water** | **Sodium hydroxide** | **Heat** |
| --- | --- | --- | --- |
| CIP | 33,990L | 113.3kg | 3,291,778 kJ |
| SIP | 1,030L |  | 117,074 kJ |

**S.4. Details of the raw materials of the basal culture medium.**

Table S.6 shows the breakdown of amino acids, glucose, vitamins, inorganic salts, and trace elements in the inventory data in Table 4 of this study.

Table S. 6. Inventory data of basal culture medium raw materials.^12^

| **Classification** | **Material** | **Current Scale** | | **Future Scale** | |
| --- | --- | --- | --- | --- | --- |
|  |  | **Amounts used/1L culture medium production** | | **Amounts used/1L culture medium production** | |
| **Amino acids** | L-Arginine | 7.15E+01 | mg | 7.13E+01 | mg |
|  | L-Cystine | 4.97E+01 | mg | 4.96E+01 | mg |
|  | L-Glutamine | 6.01E+02 | mg | 5.99E+02 | mg |
|  | Glycine | 3.07E+01 | mg | 3.06E+01 | mg |
|  | L-Histidine | 3.20E+01 | mg | 3.19E+01 | mg |
|  | L-Isoleucine | 1.08E+02 | mg | 1.08E+02 | mg |
|  | L-Leucine | 1.08E+02 | mg | 1.08E+02 | mg |
|  | L-Lysine・HCl | 1.50E+02 | mg | 1.50E+02 | mg |
|  | L-Methionine | 3.07E+01 | mg | 3.06E+01 | mg |
|  | L-Phenylalanine | 6.76E+01 | mg | 6.74E+01 | mg |
|  | L-Serine | 4.32E+01 | mg | 4.31E+01 | mg |
|  | L-Threonine | 9.77E+01 | mg | 9.75E+01 | mg |
|  | L-Tryptophane | 1.65E+01 | mg | 1.64E+01 | mg |
|  | L-Tyrosine | 7.36E+01 | mg | 7.34E+01 | mg |
|  | L-Valine | 9.67E+01 | mg | 9.64E+01 | mg |
| **Glucose** | D-Glucose monohydrate | 5.04E+03 | mg | 5.03E+03 | mg |
| **Vitamins ＋ Inorganic salts + Trace elements** | CaCl_2_・0-2H_2_O | 3.82E+02 | mg | 3.80E+02 | mg |
|  | KCl | 4.07E+02 | mg | 4.06E+02 | mg |
|  | MgSO_4_・7H_2_O | 2.04E+02 | mg | 2.03E+02 | mg |
|  | NaCl | 5.55E+03 | mg | 5.54E+03 | mg |
|  | NaHCO_3_ | 3.77E+03 | mg | 3.76E+03 | mg |
|  | NaH_2_PO_4_・2H_2_O | 1.45E+02 | mg | 1.45E+02 | mg |
|  | FeCl_3_・6H_2_O | 7.06E-02 | mg | 7.04E-02 | mg |
|  | Ca Pantothenate | 4.16E+00 | mg | 4.15E+00 | mg |
|  | α-GPC | 7.57E+00 | mg | 7.55E+00 | mg |
|  | Folic acid | 4.11E+00 | mg | 4.10E+00 | mg |
|  | i-Inositol | 7.48E+00 | mg | 7.46E+00 | mg |
|  | Niacinamide | 4.07E+00 | mg | 4.06E+00 | mg |
|  | Pyridoxine・HCl | 4.11E+00 | mg | 4.10E+00 | mg |
|  | Riboflavin | 4.13E-01 | mg | 4.12E-01 | mg |
|  | Thiamine・HCl | 4.11E+00 | mg | 4.10E+00 | mg |

**S.5. Detailed life cycle inventory data**

Table S. 7 shows the inputs and their raw materials for the target culture medium production, amounts of each input for 1 L culture medium production, and the environmental burden intensity from the LCI database IDEA Ver. 3.3 (IDEA) used and the alternative methods.^13^ Alternative methods are given below for case data on the environmental burden intensity corresponding to materials that could not be obtained from IDEA for each substance. However, only the names of the products used are given, as the specific environmental burden intensity breakdown is prohibited from disclosure under IDEA's terms of reference.

Table S. 7. Detailed life cycle inventory data.

| **Input** | **Material** | **Current Scale** | **Future Scale** | **IDEA environmental burden intensity** | **Alternative method** |
| --- | --- | --- | --- | --- | --- |
|  |  | **Amounts/1L culture medium production** | |  |  |
| Electricity | Current Electricity Mix, | 3.53E+00 kWh | 7.94E-01 kWh | electricity, Japan, FY2020 | - |
|  | Electricity Mix in 2030 | - | 7.94E-01 kWh | electricity, by energy source, LPG, Japan, FY2020 | Details are given in 2.3 Scenario Analysis in the main text. |
|  |  |  |  | electricity, by energy source, coal, Japan, FY2020 |  |
|  |  |  |  | electricity, by energy source, LNG, Japan, FY2020 |  |
|  |  |  |  | electricity, by energy source, nuclear, Japan, FY2020 |  |
|  |  |  |  | electricity, by energy source, solar, Japan, FY2020 |  |
|  |  |  |  | electricity, by energy source, wind, Japan, FY2020 |  |
|  |  |  |  | electricity, by energy source, geothermal, Japan, FY2020 |  |
|  |  |  |  | electricity, by energy source, hydro power, Japan, FY2020 |  |
|  |  |  |  | electricity, by energy source, biomass, Japan, FY2020 |  |
|  | Renewable Electricity Mix | - | 7.94E-01 kWh | electricity, by energy source, solar, Japan, FY2020 |  |
|  |  |  |  | electricity, by energy source, wind, Japan, FY2020 |  |
| Heat | - | - | 1.57E-01 MJ | energy, town gas 13A combustion | - |
| Sterilization filter | Polysulfone | 9.93E-03 g | 4.90E-03 g | miscellaneous plastics | S.5.1 Expendable |
| Pipette | Polystyrene | 4.80E+01 g | - | polystyrene |  |
| Tube | Polypropylene | 9.33E+00 g | - | polypropylene |  |
| 100mm dish | Polystyrene | 1.10E+01 g | - | polystyrene |  |
| Reactor components | Silicone rubber | 3.35E+01 g | - | silicone rubber, compound |  |
|  | ThermoPlastic Elastomer | 1.33E+01 g | - | miscellaneous plastics |  |
| Paper | Pulp | 9.00E+00 g | - | pulp, 4-digit |  |
| Plastic tweezers | Polystyrene | 4.80E+01 g | - | polystyrene |  |
| Industrial waste, Waste plastics | | 1.50E+02 g | 1.96E+00 g | incineration, industrial waste | - |
| Industrial waste, Plants and animals residue | | 1.27E+02 g | - | incineration, industrial waste, plants and animals residue | - |
| Industrial waste, Waste animal oil and waste vegetal oil | | 6.70E+01 g | - | incineration, industrial waste, waste animal oil and waste vegetal oil | - |
| Industrial waste, Waste alkali | | 7.44E+01 g | - | industrial waste treatment, waste alkali | - |
| Industrial waste, Waste paper | | 1.80E+01 g | - | incineration, industrial waste, waste paper | - |
| Sewage treatment | | 1.00E+01 L | 2.73 L | sewerage treatment | - |
| Other wastes | | 7.83E+00 g | - | incineration treatment, industrial waste, animal carcass | - |
| L-Arginine | Amino acids | 7.15E+01 mg | 7.13E+01 mg | edible amino acid | S.5.2 Amino acids |
| L-Cystine |  | 4.97E+01 mg | 4.96E+01 mg |  |  |
| L-Glutamine |  | 6.01E+02 mg | 5.99E+02 mg |  |  |
| Glycine |  | 3.07E+01 mg | 3.06E+01 mg |  |  |
| L-Histidine |  | 3.20E+01 mg | 3.19E+01 mg |  |  |
| L-Isoleucine |  | 1.08E+02 mg | 1.08E+02 mg |  |  |
| L-Leucine |  | 1.08E+02 mg | 1.08E+02 mg |  |  |
| L-Lysine・HCl |  | 1.50E+02 mg | 1.50E+02 mg |  |  |
| L-Methionine |  | 3.07E+01 mg | 3.06E+01 mg |  |  |
| L-Phenylalanine |  | 6.76E+01 mg | 6.74E+01 mg |  |  |
| L-Serine |  | 4.32E+01 mg | 4.31E+01 mg |  |  |
| L-Threonine |  | 9.77E+01 mg | 9.75E+01 mg |  |  |
| L-Tryptophane |  | 1.65E+01 mg | 1.64E+01 mg |  |  |
| L-Tyrosine |  | 7.36E+01 mg | 7.34E+01 mg |  |  |
| L-Valine |  | 9.67E+01 mg | 9.64E+01 mg |  |  |
| D-Glucose monohydrate | Glucose | 4.96E+03 mg | 4.90E+03 mg | glucose | - |
| CaCl_2_・0-2H_2_O | Inorganic salts | 3.82E+02 mg | 3.80E+02 mg | calcium chloride, dihydrate | - |
| KCl | Inorganic salts | 4.07E+02 mg | 4.06E+02 mg | potassium chloride, GLO | - |
| MgSO_4_・7H_2_O | Inorganic salts | 2.04E+02 mg | 2.03E+02 mg | magnesium chloride, hexahydrate | - |
| NaCl | Inorganic salts | 5.55E+03 mg | 5.54E+03 mg | edible salt, rock salt, solution mining method | - |
| NaHCO_3_ | Inorganic salts | 3.77E+03 mg | 3.76E+03 mg | sodium hydrogen carbonate (sodium bicarbonate) | - |
| NaH2PO_4_・2H_2_O | Inorganic salts | 1.45E+02 mg | 1.45E+02 mg | sodium phosphate | - |
| FeCl_3_・6H_2_O | Trace elements | 7.06E-02 mg | 7.04E-02 mg | ferric chloride, 38% aqueous solution | - |
| Ca Pantothenate | Vitamin | 4.16E+00 mg | 4.15E+00 mg | monosodium glutamate | S.5.2 Vitamin |
| α-GPC |  | 7.57E+00 mg | 7.55E+00 mg |  |  |
| Folic acid |  | 4.11E+00 mg | 4.10E+00 mg |  |  |
| i-Inositol |  | 7.48E+00 mg | 7.46E+00 mg |  |  |
| Niacinamide |  | 4.07E+00 mg | 4.06E+00 mg |  |  |
| Pyridoxine・HCl |  | 4.11E+00 mg | 4.10E+00 mg |  |  |
| Riboflavin |  | 4.13E-01 mg | 4.12E-01 mg |  |  |
| Thiamine・HCl |  | 4.11E+00 mg | 4.10E+00 mg |  |  |
| Egg | - | 1.05E+02 g | - | hen egg , 4-digit |  |
| Egg yolk | - | 4.07E+01 g | 1.20E+01 g | hen egg , 4-digit | Environmental burden intensity data for hen egg were distributed based on a 30% by weight ratio of yolk in egg.^14^ |
| Tap water | - | 9.76E+00 L | 1.78E+00 L | consumption, tap water | - |
|  | - |  |  | tap water | - |
| Sterilized water | Current Electricity Mix, | 0.58 kWh/m^3^ | 0.58 kWh/m^3^ | electricity, Japan, FY2020 | The value for sterilized water was calculated by adding the electricity use of 0.58 kWh/m^3^ of the sterilized water production machine and the environmental burden intensity of tap water use and consumption. |
|  | Electricity Mix in 2030 | - |  | Electricity Mix in 2030 |  |
|  | Renewable Electricity Mix | - |  | Renewable Electricity Mix |  |
|  | - | 1.36E+00 L | 2.00E+00 L | consumption, tap water |  |
|  | - |  |  | tap water |  |
| Ethanol | Ethyl alcohol | 1.93E+01 g | - | ethyl alcohol, 95% conversion | - |
| Chlorine bleach | Sodium hypochlorite | 9.19E-01 g | - | sodium hypochlorite, 12% aqueous solution | - |
| Ethylene oxide | - | 1.67E+01 g | - | ethylene oxide | - |
| Scaffold | Poly Ethylene Terephthalate filament | 2.50E+00 g | 4.80E-01 g | polyester short fiber | - |
| Salt | Salt | 1.29E-01 g | 1.40E-01 g | industrial salt | - |
| Yeast extract | - | 1.50E-01 g | 1.50E-01 g | ammonium chloride | - |
| Alkaline cleaning solution | Sodium hydroxide | - | 4.53E+00 g | sodium hydroxide | - |

S.5.1 Expendable

For the expendables used in this study, such as laboratory equipment and device consumables, detailed data for each type of expendable product were not obtained using IDEA. Therefore, this study used IDEA environmental burden intensity data based on the raw material of each expendable product.

S.5.2 Amino acids

For amino acids, the basal culture medium raw materials and detailed data for each type of amino acid were not obtained in IDEA. Therefore, IDEA environmental burden intensity data for “edible amino acids” were used for all amino acids.

S.5.3 Vitamin

Data related to vitamins, and the raw materials of the basal medium, were not obtained using IDEA. Most vitamins are industrially produced by biotechnology using fermentation methods, and the vitamin production process is similar to other fermentation processes.^15,16^ Monosodium glutamate, a type of amino acid, is also industrially produced using fermentation.^17^ Therefore, IDEA environmental burden intensity data of “monosodium glutamate” were used for all vitamins. The purification process was also considered in the intensity data.

**S.6. Life Cycle Impact Assessment (LCIA)**

This study used the Life-cycle Impact assessment Method based on Endpoint modeling 2 (LIME2), an LCIA methodology that reflects the environmental conditions of Japan in the LCIA.^18^ LIME2 finally integrates environmental impacts by a single index through characterization and damage assessment.^8^ In the characterization, the impact of the environmental load substance on each impact category was evaluated. The following 13 impact categories were evaluated: climate change, ozone layer depletion, acidification, urban area air pollution, photochemical ozone, toxic chemicals (cancer), toxic chemicals (chronic disease), aquatic toxicity, biological toxicity, eutrophication, land use (occupation), land use (transformation), and resource consumption. Damage assessment was used to evaluate the impact on 13 impact categories based on four endpoints: human health, social assets, biodiversity, and primary production. Finally, the damage assessment results for each endpoint were integrated into a single index with a monetary unit (Japanese yen, JPY) by weighting the importance among those four endpoints. This study revealed the impacts of climate change, ozone layer depletion, acidification, eutrophication, land use (occupation), land use (transformation), resource consumption on characterization, and water resources consumption. Table S.8 shows the results of the 13 impact categories on the characterization, damage assessment, integration, and water resources consumption at [Current Scale], [Future Scale,2020 mix], [Future Scale, 2030 mix], and [Future Scale, RE mix].

Table S. 8. Characterization, damage assessment, integration, and water resources consumption results.

|  | | | **Current Scale** | **Future Scale, 2020 mix** | **Future Scale,**  **2030 mix** | **Future Scale,**  **RE mix** |
| --- | --- | --- | --- | --- | --- | --- |
| **Characterization** | **Climate change** | kg-CO_2_eq | 3.32E+00 | 4.93E-01 | 3.20E-01 | 8.72E-02 |
|  | **Ozone layer depletion** | kg-CFC-11eq | 1.80E-07 | 4.96E-08 | 2.64E-08 | 2.75E-08 |
|  | **Acidification** | kg-SO_2_eq | 3.38E-03 | 4.66E-04 | 3.16E-04 | 7.73E-05 |
|  | **Urban area air pollution** | kg-SO_2_eq | 2.40E-03 | 3.49E-04 | 2.36E-04 | 5.14E-05 |
|  | **Photochemical ozone** | kg-C_2_H_4_eq | 5.48E-05 | 8.36E-06 | 5.44E-06 | 1.43E-06 |
|  | **Toxic chemicals (cancer)** | kg-C_6_H_6_eq | 1.45E-03 | 1.92E-04 | 1.91E-04 | 2.06E-04 |
|  | **Toxic chemicals**  **(chronic disease)** | kg-C_6_H_6_eq | 1.61E-05 | 2.41E-06 | 2.05E-06 | 1.67E-06 |
|  | **Aquatic toxicity** | kg-C_6_H_6_eq | 6.60E-03 | 1.32E-03 | 1.21E-03 | 1.05E-03 |
|  | **Biological toxicity** | kg-C_6_H_6_eq | 1.24E-01 | 4.72E-02 | 4.65E-02 | 4.60E-02 |
|  | **Eutrophication** | kg-PO_4_^3-^eq | 9.61E-03 | 1.11E-03 | 1.11E-03 | 1.11E-03 |
|  | **Land use (Occupation)** | m^2^a | 2.69E-01 | 3.49E-02 | 3.76E-02 | 6.29E-02 |
|  | **Land use (Transformation)** | m^2^ | 4.34E-03 | 6.86E-04 | 7.73E-04 | 1.63E-03 |
|  | **Resource Consumption** | kg-Sbeq | 3.72E-04 | 3.89E-06 | 6.20E-06 | 7.89E-06 |
| **Damage assessment** | **Human health** | DALY | 8.12E-07 | 1.19E-07 | 7.89E-08 | 2.01E-08 |
|  | **Social assets** | JPY | 6.75E+00 | 7.34E-01 | 6.16E-01 | 4.70E-01 |
|  | **Biodiversity** | EINES | 1.04E-11 | 6.36E-12 | 6.38E-12 | 6.57E-12 |
|  | **Primary production** | kg-DW | 1.21E-01 | 1.89E-02 | 3.00E-02 | 1.41E-01 |
| **Integration** | **Single index** | JPY | 1.72E+02 | 9.34E+01 | 9.34E+01 | 1.00E+02 |
| **Other** | **Water resources consumption** | m^3^ | 3.30E-02 | 4.74E-03 | 4.72E-03 | 4.71E-03 |
